# MEG/EEG group study with MNE: recommendations, quality assessments and best practices

**DOI:** 10.1101/240044

**Authors:** Mainak Jas, Eric Larson, Denis Engemann, Jaakko Leppäkangas, Samu Taulu, Matti Hämäläinen, Alexandre Gramfort

## Abstract

Cognitive neuroscience questions are commonly tested with experiments that involve a cohort of subjects. The cohort can consist of a handful of subjects for small studies to hundreds or thousands of subjects in open datasets.

While there exist various online resources to get started with the analysis of magnetoencephalography (MEG) or electroencephalography (EEG) data, such educational materials are usually restricted to the analysis of a single subject. This is in part because data from larger group studies are harder to share, but also analyses of such data are often require subject-specific decisions which are hard to document.

This work presents the results obtained by the reanalysis of an open dataset from Wakeman and Henson (2015) using the MNE software package. The analysis covers preprocessing steps, quality assurance steps, sensor space analysis of evoked responses, source localization, and statistics in both sensor and source space. Results with possible alternative strategies are presented and discussed at different stages such as the use of high-pass filtering versus baseline correction, tSSS versus SSS, the use of a minimum norm inverse versus LCMV beamformer, and the use of univariate or multivariate statistics. This aims to provide a comparative study of different stages of M/EEG analysis pipeline on the same dataset, with open access to all of the scripts necessary to reproduce this analysis.

## 1 Overview

Magnetoencephalography and electroencephalography (M/EEG) are neuroimaging technologies with a high temporal resolution, which provide non-invasive access to population-level neuronal dynamics on virtually any temporal scale currently considered relevant to cognition. While MEG can recover spatial patterns at a higher signal-to-noise ratio (SNR) and enjoys a more selective cortical resolution than EEG (Baillet, 2017), EEG is more portable and less expensive, and thus supports the study of cognition in a wider range of situations. Processing M/EEG recordings, however, is inherently challenging due to the multi-dimensional nature of the data, the low SNR of brain-related M/EEG signals, and the differences in sensitivity of these measurement techniques. This can give rise to complex sequences of data processing steps which demand a high degree of organization from the investigator.

In an effort to address reproducibility issues recently shown to affect neuroimaging studies (Ioannidis, 2005a; Button et al., 2013; Carp, 2012a,b), a number of community-led efforts have begun developing data sharing (Poldrack and Gorgolewski, 2017) and data organization (Gorgolewski et al., 2016; Galan et al., 2017) projects. These efforts are necessary first steps, but are not sufficient to solve the problem—they must be complemented by educational tools and guidelines that establish good practices for M/EEG analysis (Gross et al., 2013). However, putting guidelines into practice is not always straightforward, as researchers in the M/EEG community rely on several software packages (Tadel et al., 2011; Delorme and Makeig, 2004; Delorme et al., 2011; Oostenveld et al., 2011; Dalal et al., 2011; Litvak et al., 2011), each of which is different. Even though these packages provide tutorials for single subject data analysis, it is typically left up to the investigator to coordinate and implement multi-subject analyses. Here, we try to address this gap by demonstrating a principled approach to the assembly of group analysis pipelines with publicly available code^1^ and extensive documentation.

As members and maintainers within the MNE community, we will present analyses that make use of the MNE software suite (Gramfort et al., 2014). Historically, MNE was designed to calculate minimum-norm estimates from M/EEG data, and consisted in a collection of C-routines interfaced through bash shell scripts. Today, the MNE software has been reimplemented in (Gramfort et al., 2013a) and transformed into a general purpose toolbox for processing electrophysiology data. Built on top of a rich scientific ecosystem that is open source and free, MNE now offers state-of-the-art inverse solvers and tools for preprocessing, time-frequency analysis, machine learning (decoding and encoding), connectivity analysis, statistics, and advanced data visualization. MNE, moreover, has become a hub for researchers who use it as a platform to collaboratively develop novel methods or implement and disseminate the latest algorithms from the M/EEG community (Engemann and Gramfort, 2015; Smith and Kutas, 2015a,b; Haufe et al., 2014; King and Dehaene, 2014; Gramfort et al., 2013b; Schurger et al., 2013; Khan and Cohen, 2013; Larson and Lee, 2013; Hauk et al., 2011; Gramfort et al., 2010; Rivet et al., 2009; Kriegeskorte et al., 2008; Maris and Oostenveld, 2007). With this work, we not only share best practices to facilitate reproducibility, but also present these latest advances in the MNE community which enable automation and quality assessment.

Here, we demonstrate how to use MNE to reanalyze the OpenfMRI dataset ds000117 by Wakeman and Henson (2015). This requires setting the objectives for the data analysis, breaking them down into separate steps and taking a series of decisions on how to handle the data at each of those steps. While there may be several interesting scientific questions that have not yet been addressed on this dataset, here we confine ourselves to the analysis of well-studied time-locked event-related M/EEG components, i.e, event-related fields (ERF) and event-related potentials (ERP). This is motivated by educational purposes to help facilitate comparisons between software packages and address reproducibility concerns. To this end, we will lay out all essential steps from single subject raw M/EEG recordings to group level statistics. Importantly, we will highlight the essential options, motivate our choices and point out important quality control objectives to evaluate the success of the analysis at every step.

We will first analyze the data in sensor space. We will discuss the best practices for selecting filter parameters, marking bad data segments, suppressing artifacts, epoching data into time windows of interest, averaging, and doing baseline correction. Next, we turn our attention to source localization: the various steps involved in the process starting from defining a head conductivity model, source space, coregistration of coordinate frames, data whitening, lead field computation, inverse solvers, and transformation of source-space data to a common space. Along the way, we will present various diagnostic visualization techniques that assist quality control at each processing step, such as channel-wise power spectral density (PSD), butterfly plots with spatial colors to facilitate readability, topographic maps, and whitening plots. Finally, we will attempt to distill from our analysis, guiding principles that should facilitate successfully designing *other* reproducible analyses rather than blindly copying the recipes presented here.

## 2 Preliminaries

In this work, we describe a full pipeline using MNE to analyze the OpenfMRI dataset ds000117 by Wakeman and Henson (2015). The data consist of simultaneous M/EEG recordings from 19 healthy participants performing a visual recognition task. Subjects were presented images of famous, unfamiliar and scrambled faces. The dataset provides a rich context to study different neuroscientific and cognitive questions, such as: Which brain dynamics are characteristic of recognizing familiar as compared to unfamiliar faces? How do commonly studied face-responsive brain regions such as the Superior Temporal Sulcus (STS), the Fusiform Face Area (FFA) and the Occipital Face Area (OFA) interact when processing the familiarity of the face? At the same time, it presents a well-studied paradigm which can be particularly beneficial for the development of methods related to connectivity and source localization.

### 2.1 Data description

The subjects participated in 6 runs, each 7.5 minutes in duration. In the original study, three subjects were discarded due to excessive artifacts in the data. To produce comparable results, the same subjects are also discarded from the group results in this study. The data were acquired with an Elekta Neuromag Vectorview 306 system consisting of 102 magnetometers and 204 planar gradiometers. In addition, a 70 channel Easycap EEG system was used for recording EEG data simultaneously.

### 2.2 Reading data

MNE supports multiple file formats written by M/EEG hardware vendors. Apart from Neuromag *FIF* files, which are the default storage format, MNE can natively read multiple other formats ranging for MEG data including 4D Neuroimaging BTI, KIT, and CTF, and for EEG data B/EDF, EGI, and EEGLAB *set*^2^. Despite this heterogeneity of systems, MNE offers a coherent interface to the metadata of the recordings using the so-called *measurement info*^3^. Regardless of the input format, all processed files can be saved as *FIF* files or in the HDF5 format^4^.

MNE can handle multimodal data containing different channel types, the most common being magnetometer, gradiometer, EEG, electrooculogram (EOG), electrocardiogram (ECG), and stimulus trigger channels that encode the stimulation paradigm. MNE also supports electromyogram (EMG), stereotactic EEG (sEEG) and electrocorticography (ECoG), functional near-infrared spectroscopy (fNIRS) or miscellaneous (misc) channel types. Declaring and renaming channel types is a common step in the preparation of M/EEG datasets before analysis. In our case, once the files were read in, some of the channels needed to be renamed and their channel types corrected in the measurement info (see (Wakeman and Henson, 2015)): the EEG061 and EEG062 electrodes were set as EOG, EEG063 was set as ECG, and EEG064 was set as a miscellaneous channel type as it was a free-floating electrode. If this step is omitted, some preprocessing functions may fall back to potentially less optimal defaults, for example, using the average of the magnetometers instead of the ECG channel when searching for cardiac events.

## 3 MEG and EEG data preprocessing

### 3.1 Maxwell filtering (SSS)

Neuromag MEG recordings are often preprocessed first using the Signal Space Separation (SSS) method (Taulu, 2006), otherwise known as Maxwell filtering. SSS decomposes the data using multipole moments based on spherical harmonics and removes the component of magnetic field originating from outside the MEG helmet. SSS is therefore useful for removing environmental artifacts, and can also be used to compensate for head movements during the recording. In this study, movement compensation is not strictly necessary as the participants managed to stay predominantly still.

The data provided by OpenfMRI (Poldrack and Gorgolewski, 2017) already contain files processed using the proprietary Elekta software MaxFilter, which is what we use in our analysis for the sake of reproducibility. However, MNE offers an open source reimplementation and extension of SSS as well. Before running SSS, it is crucial that bad channels are marked, as otherwise SSS may spread the artifacts from the bad channels to all other MEG channels in the data. This step is preferably done manually with visual inspection. When using the MNE implementation of Maxwell filtering, we reused the list of bad channels available from the Elekta MaxFilter logs in the dataset.

Results comparing raw data, data processed by Elekta MaxFilter, and data processed by the MNE `maxwell_filter` function are provided in Figure 1. While the unprocessed data do not show a clear evoked response, the Maxwell filtered data do exhibit clear event-related fields with a clear peak around 100 ms post-stimulus. Note that the results obtained with Elekta implementation and the MNE implementation have minimal differences due to slight differences in computation of component regularization parameters.

**Figure 1:**
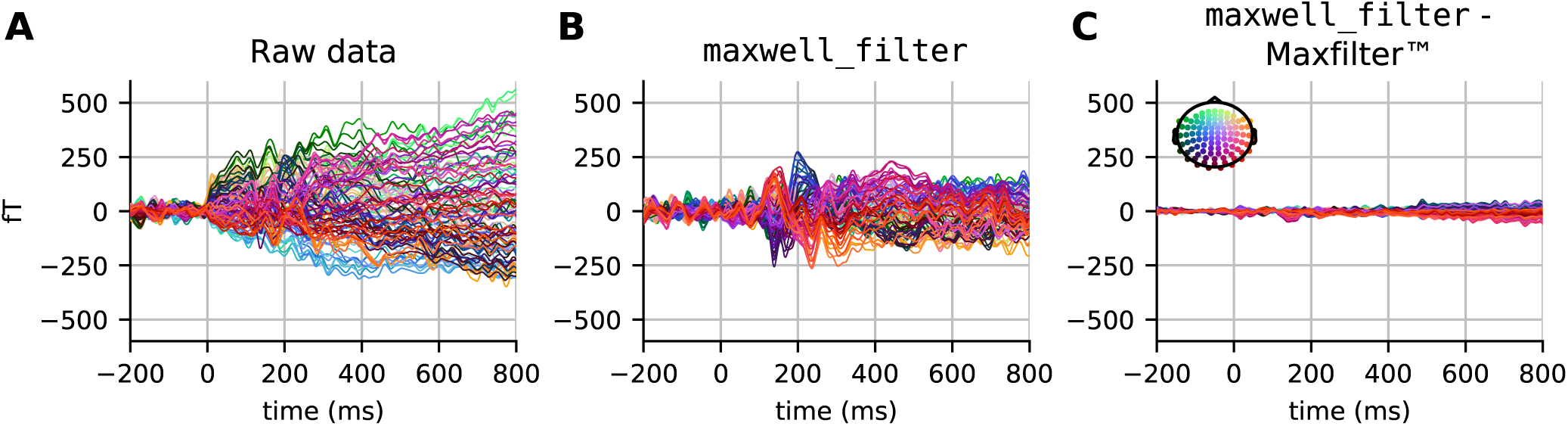
Evoked responses (filtered between 1 and 40 Hz) in the magnetometer channels from (A) unprocessed data, (B) data processed with `maxwell_filter` in MNE, and (C) the difference between data processed using `maxwell_filter` and Elekta MaxFilter (TM). The colors show the sensor position, with *(x,y,z)* sensor coordinates converted to *(R,G,B)* values, respectively.

#### Alternatives

In principle, SSS can be applied to data acquired with any MEG system providing it has comprehensive sampling (more than about 150 channels). However, so far it has not been tested extensively with other than the 306-channel Neuromag systems. SSS requires relatively high calibration accuracy, and the Neuromag systems are thus carefully calibrated for this purpose. If SSS is not an option, for example due to the lack of fine-calibration information, reasonable noise reduction can be readily obtained from Signal Space Projections (SSP) (Uusitalo and Ilmoniemi, 1997). This intuitively amounts to projecting out spatial patterns of the empty room data covariance matrix using Principal Component Analysis (PCA). In practice, depending on the shielding of the room, up to a dozen SSP vectors can be discarded to obtain satisfactory denoising.

#### Caveats

It is important to highlight that after SSS, the magnetometer and gradiometer data are projected from a common lower dimensional SSS coordinate system that typically spans between 64 and 80 dimensions. As a result, both sensor types contain highly similar information, which also modifies the inter-channel correlation structure. This is the reason why MNE will treat them as a single sensor type in many of the analyses that follow.

### 3.2 Power spectral density (PSD)

The power spectral density (PSD) estimates for all available data channels provide a convenient way to check for spectral artifacts and, in some cases, bad channels. MNE computes the PSD of raw data using the standard Welch’s method (Welch, 1967; Percival and Walden, 1993), whereby the signal for each channel is analyzed over consecutive time segments, with eventually some overlap. Each segment is windowed and then the power of the discrete Fourier transform (DFT) coefficients is computed and averaged over all segments. By making the assumption that each of these segments provides a realization of a stationary process, the averaging procedure produces an unbiased estimate of the PSD with reduced noise.

Starting from MNE version 0.14, we show channel-wise PSD plots rather than an average across channels, as this facilitates spotting outlier channels. In Figure 2, we show the PSD for the EEG channels in one run for one subject. We use windows of length 8192 samples (about 7.4 s given the 1.1 kHz sampling rate) with no overlap. Using a power of 2 for the length and no overlap accelerates computations. Using a logarithmic frequency-axis scaling for the PSD enables quality control by facilitating screening for bad channels. In fact, we found that some potentially bad channels (e.g., EEG024 in subject 14 for run 01) were omitted by the authors of (Wakeman and Henson, 2015), although they are clearly visible in such plots. Concretely we see a few channels with strongly increased low-frequency power below 1 Hz. On the other hand, using a linear frequency-axis scaling, we can convince ourselves easily that the data is unfiltered, as it contains clear peaks from power line at harmonics of 50 Hz, as well as the five Head Position Indicator (HPI) coils used to monitor the head position of the subject, at frequencies of 293, 307, 314, 321, and 328 Hz.

**Figure 2:**
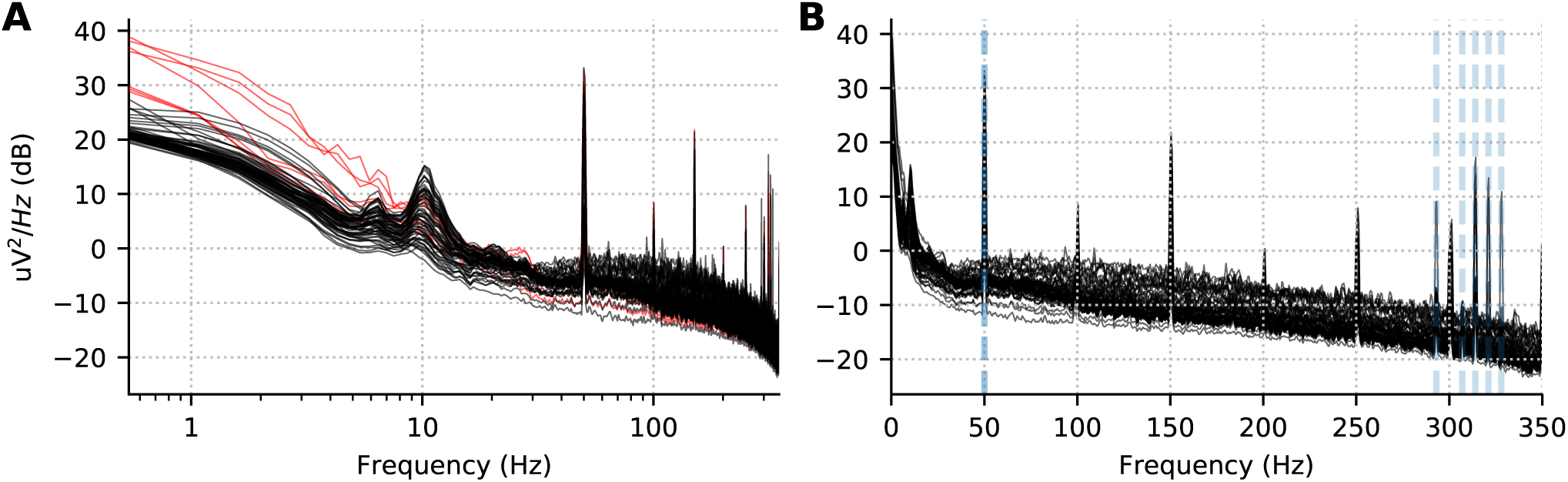
Power spectral density per channel for subject 10, run 02. (A) Log scale for the x axis accentuates low frequency drifts in the data. The red lines show the PSD for the bad channels marked manually and provided to us by Wakeman and Henson (2015). (B) The same data with a linear x-axis scale. Five peaks corresponding to HPI coils around 300 Hz are visible and marked in gray dotted lines alongside the power line frequency (50 Hz).

#### Alternatives

The same could have been achieved with the multitaper method (Percival and Walden, 1993; Slepian, 1978), where the data is multiplied element-wise by orthogonal data tapers. However, this method can be an order of magnitude slower than the Welch method for long continuous recordings. The multitaper method is indeed recommended for short data segments. Here we are interested in the PSD for diagnostic purposes on the raw continuous data, and we therefore use the Welch method, a.k.a. averaged periodogram method.

### 3.3 Temporal filtering

In this study, we focused on event-related brain signals below 40 Hz. We low-pass filtered our data at a 40 Hz cutoff frequency with 10 Hz transition band. Such a filter does not affect ERP signals of interest, attenuates the line frequency of 50 Hz and all HPI coil frequencies. It also limits the effects of temporal ringing thanks to a wide transition band. Because the low-pass was sufficiently low, we did not employ a notch filter separately. Note that such a choice of filters is not necessarily a good default for all studies of event-related brain responses, as ERFs or ERPs can contain rather high frequencies (see for example (Götz et al., 2015)).

When filtering, it is important to take into account the frequency response and impulse response of the filter. In MNE 0.16, the default filter will adapt the filter length and transition band size based on the cutoff frequencies, as done in the EEGLAB software (Widmann et al., 2015; Parks and Burrus, 1987; Ifeachor and Jervis, 2002)^5^. Although no default parameters will fit all analysis requirements, MNE chooses parameters that aim to achieve reasonable stop-band attenuation without excessive filter ringing. To illustrate this point, we compare filters across MNE versions using frequency response and impulse response plots in Figure 3. The stop-band attenuation and transition bandwidth in Figure 3A and Figure 3B are less restricted in the newer versions, which results in less steep attenuation but also less temporal ringing in the impulse response (see Figures 3C and D). It can be seen that the previous default parameters gave rise to stronger filtering artifacts as indicated by higher impulse response amplitudes across the time window.

**Figure 3:**
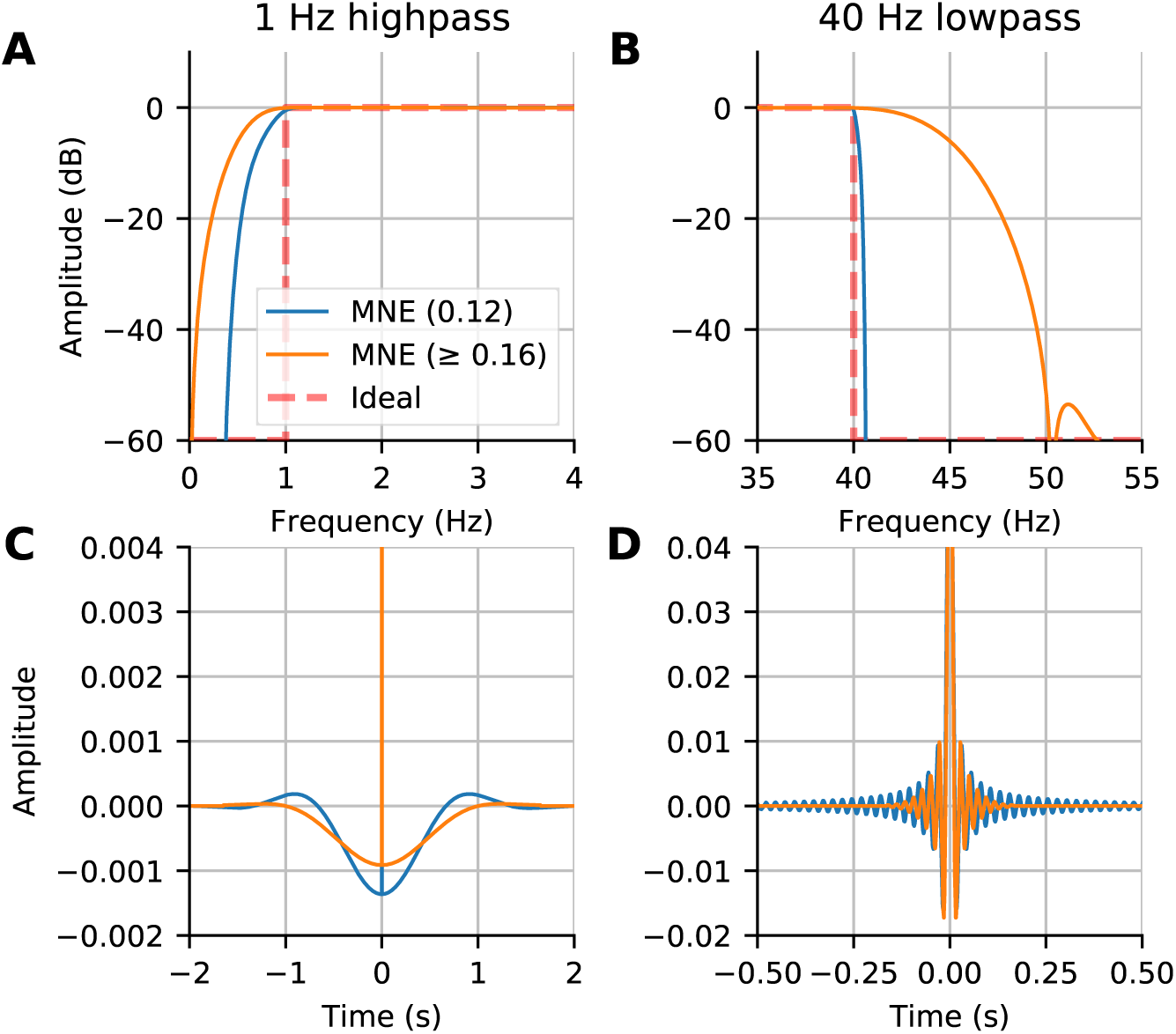
Comparison of filters between new (0.16) and old (0.12) MNE versions: (A) The frequency response of the highpass filter; (B) The frequency response of the lowpass filter; (C) The impulse response of the highpass filter; (D) The impulse response of the lowpass filter. The filters in MNE are now adaptive with trade-offs between frequency attenuation and time domain artifacts that by default adapt based on the chosen low-pass and high-pass frequencies.

#### Alternatives and Caveats

If the signal quality is satisfactory, filtering may not be necessary. In the context of this study, we decided to baseline correct our signals rather than high-pass filter them, keeping in mind the ongoing discussion in the community on this topic (Tanner et al., 2015; Rousselet, 2012; Widmann and Schröger, 2012; Acunzo et al., 2012; Maess et al., 2016). Our choice will be motivated in Section 3.7 on baseline correction.

### 3.4 Marking bad segments and channels

The next step in the pipeline is to remove bad data segments and bad channels. As data have been processed with Maxwell filter, there are no more bad MEG channel at this stage. For the bad EEG channels, we use the ones provided by the original authors.

To remove bad data segments and bad epochs due to transient artifacts, it is possible in MNE to use the epochs plotter interactively, or to do it via scripting. Either way, the indices of all epochs that are removed from further analysis are logged in the *drop log* attribute of the epochs objects (see online documentation of the Epochs class^6^).

As we are building a reproducible pipeline, here we prefer the scripting route. In MNE, this can be achieved by removing trials whose peak-to-peak amplitude exceeds a certain rejection threshold. Even though this works reasonably well for single subject analysis, it would likely need to be tuned for individual subjects in group studies. Therefore, instead of specifying the thresholds manually, we learn it from the data using the *autoreject* (global) (Jas et al., 2017) algorithm. *Autoreject* is an unsupervised algorithm which minimizes the cross-validation error, measured by the Frobenius norm between the average signal of the training set and the median signal of the validation set. *Autoreject* not only removes trials containing transient jumps in isolated MEG or EEG channels, but also eyeblink artifacts affecting groups of channels in the frontal area. Since we are dealing with visual stimuli, it is preferable to remove the eyeblink trials altogether using the EOG rejection threshold over the stimulus presentation interval rather than suppressing the artifact using a spatial filter such as ICA or SSP. Given the large number of trials at our disposal, we can afford to remove some without affecting the results very much.

For the purpose of group averaging, the bad EEG channels were repaired by spherical spline interpolation (Perrin et al., 1989) so as to have the same set of channels for each subject.

### 3.5 Independent Component Analysis (ICA)

Bad channel or segment removal can correct for spatially and temporally isolated artifacts. However, it does not work well for systematic physiological artifacts that affect multiple sensors. For this purpose, ICA is commonly used (Jung et al., 1998). ICA is a blind source separation technique that maximizes the statistical independence between the components. While PCA only requires orthogonal components, ICA looks for independence for example by looking at higher statistical moments beyond (co)variance. In the context of MEG and EEG analysis, common physiological artifacts have skewed and peaky distributions, hence are easily captured by ICA methods that look for non-Gaussian sources. ICA is therefore popular for removing eye blinks and heart beats, which manifest themselves with prototypical spatial patterns on the sensor array.

In the present study, we use FastICA (Hyvarinen, 1999) to decompose the signal into maximally independent components. We estimate the ICA decomposition on band-pass filtered (1 Hz highpass with 1 Hz transition band, 40 Hz lowpass with 10 Hz transition band) data that has been decimated. In practice, to improve the quality of ICA solution, high-pass filtering is often helpful as it can help to minimize violations of the stationarity assumption made by ICA. Likewise, it is recommended to exclude data segments containing environmental artifacts with amplitudes higher than the artifacts of interest. Finally, generous decimation can save computation time and memory without affecting the quality of the ICA solution, at least, when it comes to separating physiological artifacts from brain signals. Both measures can be implemented using the `reject` and `decim` parameters provided by the ICA fitting routine in MNE. Here we decimated the data by a factor of 11, and excluded time segments exceeding amplitude ranges of 4000 × 10^−13^ fT cm^-1^ and 4 × 10^−12^ fT on the magnetometers and gradiometers, respectively.

The ICA component corresponding to ECG activity is then identified using cross-trial phase statistics (CTPS) (Dammers et al., 2008) using the default threshold of 0.8 on the Kuiper statistic. Pearson correlations are used to find EOG related components. As ICA is a linear model, the solution can be estimated on continuous *raw* data and subsequently used to remove the bad components from the *epochs* or *evoked* data.

#### Alternatives

MNE also implements CORRMAP (Viola et al., 2009) which is particularly useful when no ECG or EOG channels are available. This approach uses pattern matching of ICA spatial components. Once templates have been manually defined for one subject, similar patterns can be found for the remaining subjects. If ICA is not an option, SSP projections provide a simple and fast alternative. Here, they can be computed from time segments contaminated by the EOG and ECG artifacts and commonly the first 1 to 2 components are projected out. In our experience, SSP is less precise in separating artifacts from brain components than ICA for the reasons mentioned above, yet, often good enough for a wide class of data analysis scenarios. For analysis of single EEG sensors, multivariate methods cannot be applied. Computing the residuals of a linear regression from the ECG sensor on the EEG is an option in this case.

#### Caveats

Before blindly applying ICA, it is recommended to estimate the amount of contamination of the MEG and EEG signals. This can be easily achieved by detecting artifact events and epoching and averaging the data accordingly. If, for example, the amplitude range of the average ECG artifact is close to the amplitude range of the brain signals and only few events occur, chances are low to estimate clear cut ECG components using ICA. However, in this case the contamination by ECG is low and therefore no advanced artifact suppression is needed. Second, there is a trade-off between processing time and accuracy. For many analyses, mitigating the artifact contamination by a significant proportion is sufficient and methods like SSP are a reasonable choice. In certain decoding analyses, such preprocessing considerations may have little relevance if any for the final classification results. Indeed, the combination of supervised and multivariate decoding algorithms allows to extract the signals of interest directly in one step.

### 3.6 Epoching

In event-related M/EEG studies, a trigger channel (in this data STI101) contains binary-coded trigger pulses to mark the onset/offset of events. These pulses can be automatically extracted from the data during analysis and the values on the trigger channel are mapped to the *event IDs*. MNE offers the possibility to extract events when the signal in the trigger channel increases, decreases, or both. It also allows the construction of binary masks to facilitate selecting only the desired events. We masked out the higher order bits in the trigger channel when extracting the events as these corresponded to key presses. After extraction, events can be freely manipulated or created as necessary by the user, as they only require i) the sample number, and ii) some integer code relevant for the experiment or analysis.

As a next step, we extracted segments of data from the continuous recording around these events and stored them as single trials, which are also called epochs, in MNE. The `Epochs` object can store data for multiple events and the user can select a subset of these as `epochs[event_id]`^7^. Moreover, MNE offers the possibility for the user to define a hierarchy of events by using tags (similar in flavor to hierarchical event descriptors by Bigdely-Shamlo et al. (2013)). This is done using `event_id` which is a dictionary of key-value pairs with keys being the tags separated by a forward slash (/) and values being the trigger codes^8^. For the paradigm used in this study we used:

1. events_id = {
2. ‘face / famous /first’:5,
3. ‘ face/famous/immediate’:6,
4. ‘ face/famous/long’:7,
5. ‘ face / un familiar / first’:13,
6. ‘ face / un familiar / immediate’:14,
7. ‘ face / un familiar / long’:15,
8. ‘ scrambled / first’:17,
9. ‘scrambled /immediate’:18,
10. ‘ scrambled / long’:19,
11. }

At the highest level of hierarchy are ‘face’ and ‘scrambled’. A ‘face’ can be ‘famous’ or ‘unfamiliar’. And a famous face can be ‘first’, ‘immediate’ or ‘long’ (This distinction between the three categories of famous faces was not used in our analysis). Later on, accessing all the epochs related to the ‘face’ condition is straightforward, as one only needs to use `epochs[‘face’]` and MNE internally pools all the sub-conditions together. Finally, the epochs were constructed starting 200 ms before stimulus onset and ending 2900 ms after (the earliest possible time of the next stimulus onset).

### 3.7 Baseline correction

It is common practice to use baseline correction so that any constant offsets in the baseline are removed. High-pass filtering achieves similar results by eliminating the low-frequency components in the data. However, when using baseline correction, the low frequency drifts present in the data are not attenuated. Thus it is useful to examine long time-courses of the data, if possible, to determine if low-frequency drifts are present. The difference between the two approaches can be seen in Figure 4. The evoked responses in the figure are across-trial averages for the famous face condition. If a maximum time of approximately one second were used, a simple baseline correction would appear to produce an undesired “*fanning*” in the later responses. Indeed one can observe in Figure 4A that at one second post-stimulus, the channels still significantly deviate from zero. However, by extending the time window much longer (here to 2.9 seconds) we can see that the signals do mostly return to the baseline level.

**Figure 4:**
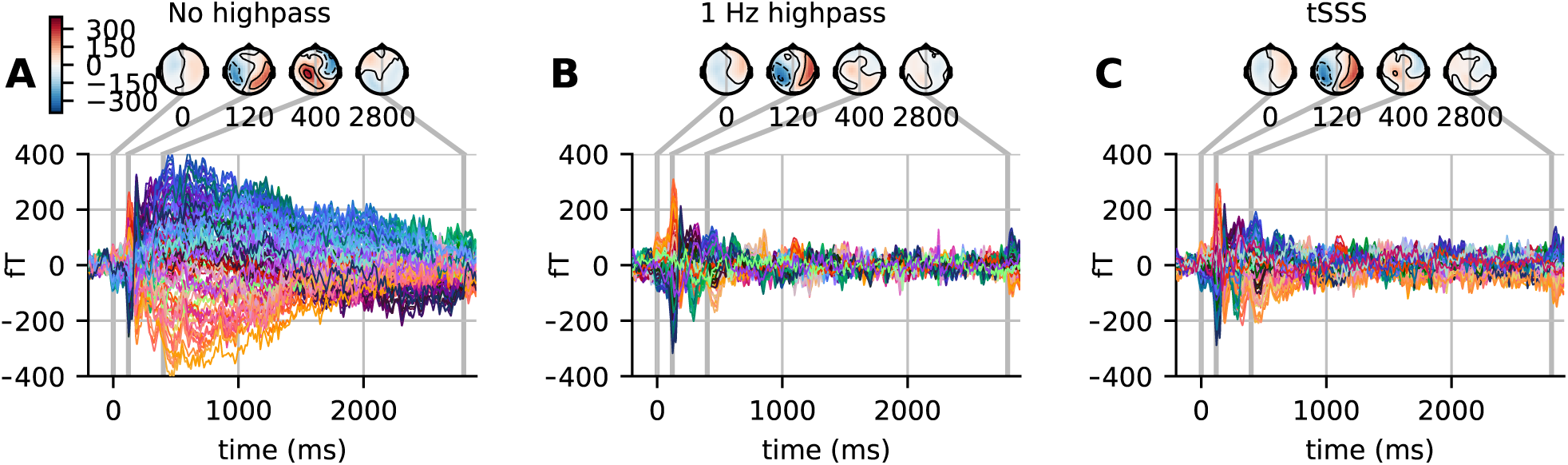
(A) Evoked response in magnetometers for subject 3 with baseline correction. Note how signals tend toward the baseline late in the epochs (where the rightmost time point, 2.9 sec, is the earliest possible start time for the next stimulus). (B) The highpass filtered version of the signal and (C) the signal processed with temporal SSS (tSSS). Both reduce the magnitude of the slow and late sustained responses shown in (A).

#### Caveats and Alternatives

With highpass filter at 1 Hz (and 1 Hz transition band), the signal returns to the baseline level much sooner. Note also the similarities between Figures 4B and 4C, illustrating how using temporal version of the SSS algorithm (tSSS) acts implicitly as a high-pass filter. For tSSS, we use a buffer size of length 1 s and a correlation limit of 0.95 to reject overlapping inner/outer signals. However, these high-passing effects come at the expense of distorting the sustained responses. We will thus focus on analyses that utilize the baseline-corrected data here.

## 4 Sensor space analysis

An important step in analyzing data at single-subject and group levels is sensor-space analysis. Here we show how several different techniques can be employed to understand the data.

### 4.1 Group average

A classical step in group studies is known as “grand averaging” (Delorme et al., 2015). It is particularly common for EEG studies and it consists in averaging ERPs across all subjects in the study. As not all subjects have generally the same good channels, this step is commonly preceded by an interpolation step to make sure data are available for all channels and for all subjects. Note that grand averaging is more common for EEG than for MEG, as MEG tends to produce more spatially resolved topographies that may not survive averaging due to signal cancellations.

The grand average of the 16 subjects for one EEG sensor (EEG065) is presented in Figure 5. We selected this channel to compare with the figure proposed by Wakeman and Henson (2015). We present the grand average for the ‘scrambled’, ‘famous’, and ‘unfamiliar’ conditions using a high-pass filter (cf. Section 3.7), and baseline corrected using prestimulus data. This figure replicates the results in (Wakeman and Henson, 2015). We can see the early difference between faces, familiar or unfamiliar, and scrambled faces around 170 ms. We can also notice a difference in the late responses between the two conditions ‘unfamiliar’ and ‘famous’. However, the effect is smaller when using high-pass filtering, as it corrects for the slow drifts.

**Figure 5:**
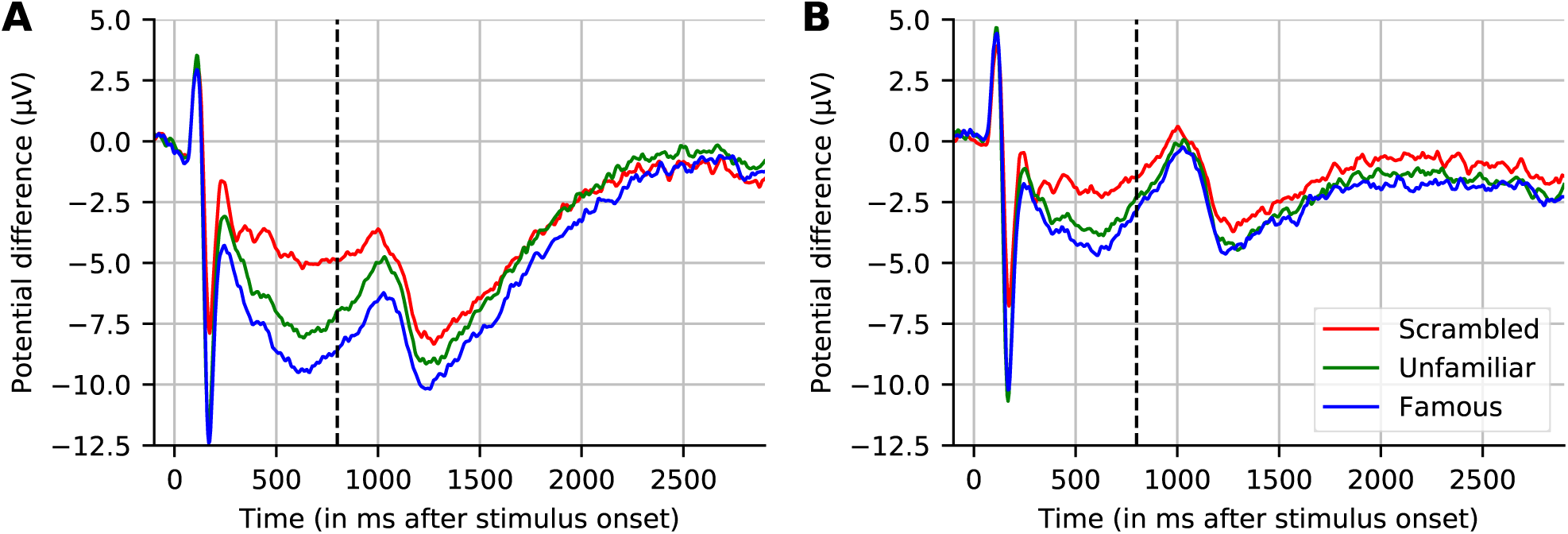
Grand averaged evoked response across 16 subjects for channel EEG065. (A) No highpass filter. (B) Highpass filtered at 1.0 Hz. Note that, similar to (A), the results reported by Wakeman and Henson (2015) (dashed line at 800 ms indicates where their plot stopped) show large drifts, but these return to near-baseline levels toward the end of a sufficiently long interval (here, 2.9 seconds) even without applying a highpass filter.

#### Caveats

For MEG, the grand average may wash out effects or induce spurious patterns due to misalignment between head positions. SSS can be used to align subjects in one common coordinate systems.

### 4.2 Contrasting conditions

Two conditions of interest are often compared using a statistical contrast. A paired contrast between two conditions can be computed by computing the difference in their evoked responses. The difference does not take into account the number of trials used to compute the evoked response - in other words, each condition is weighted equally. Recall that the event IDs were organized hierarchically during epoching (as described in Section 3.6). Such a hierarchical organization is natural for contrasting conditions in the experiment, as we compare not only ‘faces’ against ‘scrambled faces’, but also ‘famous faces’ against ‘unfamiliar faces’.

#### Caveats

Although this is standard in EEG pipelines, historically, for computing the source estimates, weighted averages have sometimes been used. However, MNE provides a mathematically correct estimate for the effective number of trials averaged, so equal-weighted combinations (additions or subtractions) of evoked data are properly accounted for even in the context of unequal trial counts. This logic, however, does not apply when working with experimental protocols (for example, oddball tasks) which, by design, produce many more examples of one than the other conditions.

### 4.3 Cluster statistics

To contrast our conditions of interest, here we use a non-parametric clustering statistical procedure as described by Maris and Oostenveld (2007). This combines neighboring values that are likely to be correlated (here, neighboring time instants) to reduce the problem of multiple comparisons. The contrast score (here the t-values) for each cluster are summed up to compute the mass of each cluster, which serves as our actual statistic. Next, we need to know if the distribution data in our two conditions (here measured using cluster sizes) is significantly different from what would be obtained by chance. For this purpose, we generate a null distribution from the data by randomly swapping our conditions for each subject according to exchangability under our null hypothesis. In this case, it is equivalent to changing the sign of the contrast data (as we are using a one-sample t-test on the difference between conditions), and then recomputing the maximal cluster size for each permutation. From an estimate of the distribution of the maximum cluster size under the null-hypothesis, we can compute the probability of observing each cluster relative to this distribution. This gives us a control of the family-wise error rate (FWER), a.k.a. type 1 error, when reporting a significant difference between the distribution of data in our two conditions.

Running this nonparametric permutation test on the single sensor EEG065 (also used by Wakeman and Henson (2015)) revealed two across-time clusters that allowed us to reject the null hypothesis at the level *p* < 0.01. To perform the clustering, we used an initial thresholding of *p* < 0.001 with a two-sided paired t-test (Figure 6). The statistic used was a one-sample t-test on the contrast ERPs using as contrast weights (0.5 for ‘familiar’, 0.5 for ‘unfamiliar’ and -1 for ‘scrambled’), testing for the condition faces versus scrambled faces. A first cluster appears around the same time as the evoked response, and the other captures the late effects. Running another statistical test, this time incorporating the spatial distribution of the sensors into the clustering procedure, yields one spatiotemporal cluster with *p <* 0.05 for the contrast condition as shown in Figure 7.

**Figure 6:**
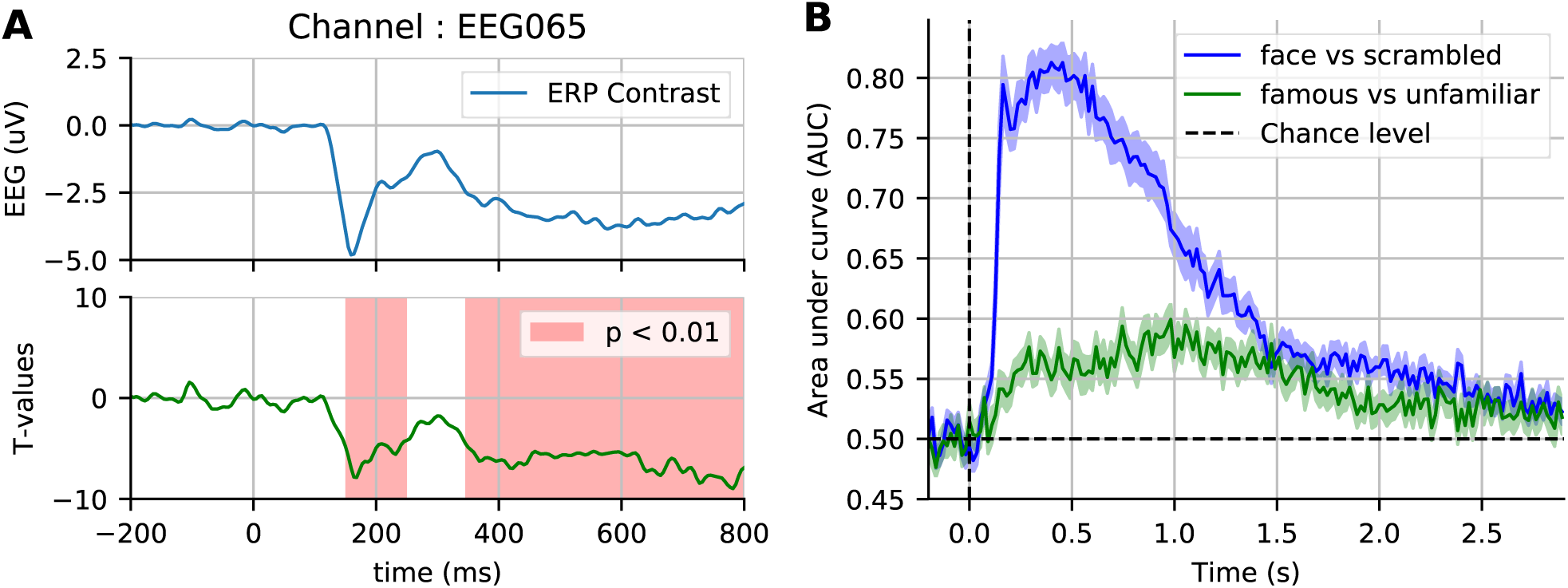
Sensor space statistics. (A) A single sensor (EEG065) with temporal clustering statistics. The clustering is based on selecting consecutive time samples that have exceeded the initial paired t-test threshold (0.001), and finding clusters that exceed the size expected by chance according to exchangability under the null hypothesis (*p <* 0.01, shaded areas). (C) Cross-validation score of time-by-time decoding. As opposed to a cluster statistic, time decoding is a multivariate method which pools together the signal from different sensors to find discriminative time points between two conditions.

**Figure 7:**
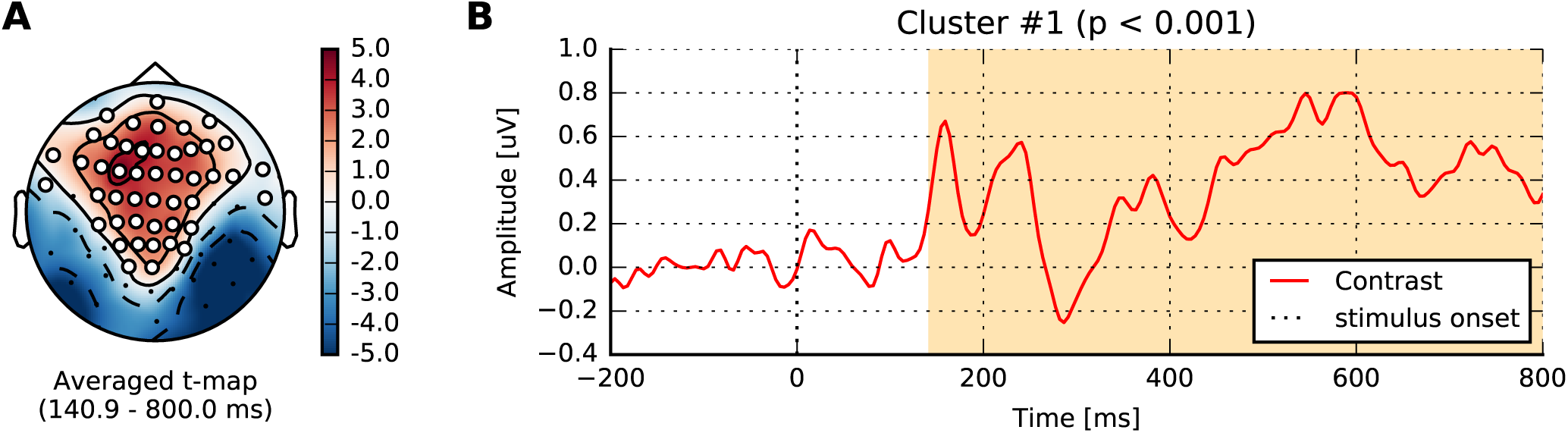
Spatiotemporal cluster statistics on the EEG sensors. (A) Topographic map of the t-statistic. (B) Average over the sensors that were part of the significant cluster.

#### Alternatives and caveats

It is important to note that this clustering permutation test does not provide feature-wise (vertex, sensor, time point, etc.) but *cluster-level* inference. This is because the test statistic is the cluster size and not any specific t-values used to obtain the cluster in the first place. When inspecting a significant cluster, no conclusion can be drawn on which time point or location was more important. A computationally more expensive alternative is the so-called threshold-free cluster enhancement (TFCE) method which provides feature-level inference and, moreover, mitigates the problem of having to set the initial threshold on the t-values to define clusters (Smith and Nichols, 2009). When strong *a priori* hypotheses exist considering few regions of interest in either time, frequency or space can be a viable alternative. In that case, the multiple comparisons problem may be readily alleviated by more conventional measures, such as false discovery rates (FDR) (Genovese et al., 2002).

### 4.4 Time Decoding

As an alternative to mass-univariate analysis, a event-related brain dynamics can studied using a multivariate decoding approach (Ramkumar et al., 2013; King and Dehaene, 2014). Here, a pattern classifier, often a linear model (e.g. logistic regression) is trained to discriminate between two conditions: ‘face’ versus ‘scrambled’, and also ‘famous faces’ versus ‘unfamiliar faces’. The classifier can be trained on single trials, time-point by time-point. The prediction success can then be assessed with cross-validation at every instant, yielding an intuitive display of the temporal evolution of discrimination success. In Figure 6B, we display such cross-validation time-series averaged across the 16 subjects. As anticipated, discriminating between faces and scrambled faces is much easier than discriminating between ‘famous’ and ‘unfamiliar’ faces, based on information in early components in the first second after stimulus-onset.

For performance evaluation, we use is area under the receiver operating characteristic curve (ROC-AUC), as it is a metric that is insensitive to class imbalance (i.e., differing numbers of trials) therefore allowing us to average across subjects, and also to compare the two classification problems (faces vs. scrambled and familiar vs. unfamiliar). Results on the faces vs. scrambled conditions show that time-resolved decoding reveals decoding accuracy greater than chance around the same time intervals as the non-parametric cluster statistic. The effect although appears here quite sustained over time. Results on familiar *vs*. unfamiliar conditions are also above chance from 200 to 300 ms, however the best decoding performance emerges later for this contrast. This more subtle effect even peaks after 800 ms, which exceeds the time window investigated in the original study.

## 5 Source reconstruction

The MNE software relies on the FreeSurfer package (Dale et al., 1999; Fischl et al., 1999) for the processing of anatomical MRI images. This automatic procedure is run using the command `recon-all` on the T1 MRI of each subject. This provides many useful pieces of information, but the most critical here are the cortical reconstruction (a high resolution triangulation of the interface between the white and the gray matter) and the inner skull surface.

For inverse source reconstruction and beamforming, we must first compute the forward solution, often called a gain or lead field matrix. It describes the sensitivity of the sensors to a given set of dipoles (Mosher et al., 1999). Computing the gain matrix, which is a linear operator, requires having a so-called source space of dipole locations, a conductor model for the head, and the sensor locations relative to those dipoles. This latter requirement in practice means putting in the same coordinate system the MRI (where the source space and conductor model are defined), the head (where the EEG electrodes are digitized), and the MEG device (where the MEG sensors are defined). This step is commonly referred to as *coregistration*. We will cover each of these steps below.

### 5.1 Source space

As we expect most of our activations of interest to be due to cortical currents (Dale et al., 2000a), we position the candidate dipoles on the cortical mantel. We chose a source space obtained by recursively subdividing the faces of an octahedron six times (oct6) for both the left and right hemispheres. This leads, for each subject, to a total of 8196 dipoles evenly spaced on the cortical surface (See Figure 6 in (Gramfort et al., 2014)).

### 5.2 Head conductivity model

MNE can use simple spherical conductor models but when the MRI of subjects are available, the recommended approach is to use a piecewise-constant conductivity model of the head. Tissue conductivities are defined for each region inside and between the segmented interfaces forming the inner skull, outer skull and the outer skin. It corresponds to a so-called three layer model, however a single layer is possible when using only MEG data. The default electrical conductivities used by MNE are 0.3 S/m for the brain and the scalp, and 0.006 S/m for the skull, i.e., the conductivity of the skull is assumed to be 1/50 of that of the brain and the scalp. With such a head model, Maxwell equations are solved with a boundary element model (BEM).

In addition to the T1 MRI image, fast low-angle shot (FLASH) images are provided in the present dataset. Such MRI images allow to automatically extract precise surfaces for the inner skull and outer skull. Note that in the absence of FLASH images, MNE offers a somewhat less accurate solution based on the watershed algorithm. One output of the MNE automatic BEM surface extraction is presented in Figure 8. It contains the three surfaces needed for the computation of the EEG gain matrix. In our results shown here, we used only the MEG data for source reconstruction, and consequently only made use of the inner skull surface in a one-layer model. As MRIs shared here are defaced, outer skull and scalp surfaces are anyway quite wrong, so we considered it satisfactory to only use the inner skull surface.

**Figure 8:**
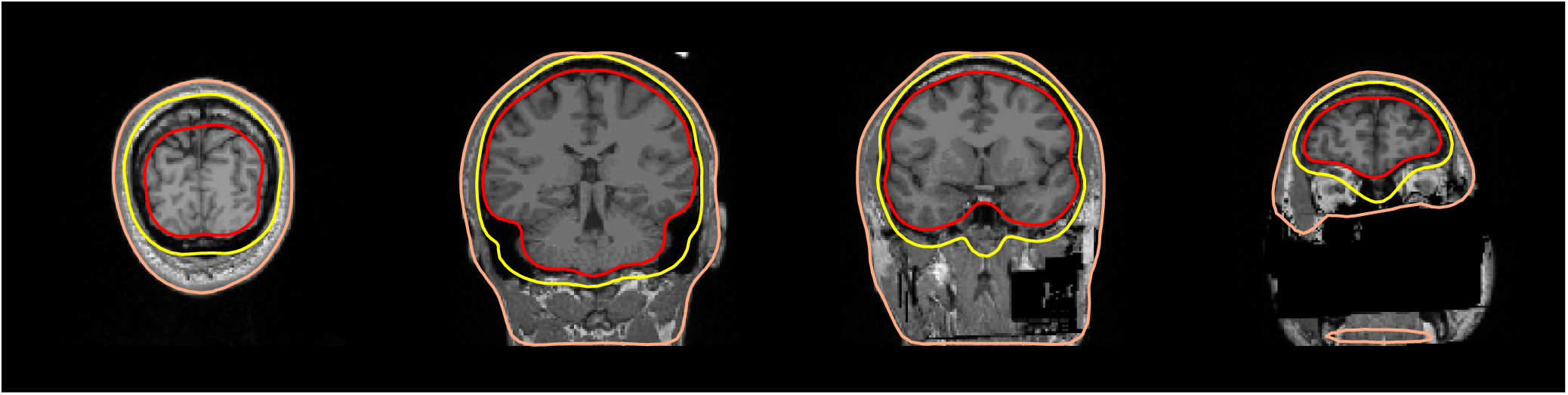
BEM surfaces on flash MRI images. The inner skull, outer skull and outer skin are outlined in color.

Quality insurance at this stage consists in checking that the three surfaces do not intersect with each other and that they follow the interfaces between the brain, the skull and the skin. A slice-by-slice visual inspection of approximate alignment is best and is conveniently proposed by MNE BEM plotting function that outputs a figure as presented in Figure 8.

Here, as the MRIs shared in this dataset were anonymized, the outer skin surface obtained automatically using Freesurfer intersected with the outer skull surface for most subjects. However this is rarely observed with non defaced T1 MRI images.

### 5.3 Coregistration

In order to compute the gain matrix, the sensor locations (and normals), head model, and source space must be defined in the same coordinate system. In practice, this means that the BEM surfaces and source space (which are defined in MRI coordinates) must be coregistered with the EEG sensors, which are digitized in the Neuromag head coordinate frame (defined by the digitized nasion, LPA, and RPA). The MEG sensor locations and normals are defined in the MEG device coordinate frame. Typically, the MEG-to-head transformation is determined during acquisition using head position indicator (HPI) coils (or redefined using head position transformation using Maxwell filtering), so MEG sensors can be easily transformed to head coordinates. The transformation between the MRI and head coordinate frames is typically estimated by identifying corresponding points in the head and MRI coordinate systems, and then aligning them.

The most common points used to provide an initial alignment are the fiducial landmarks that define the Neuromag head coordinate frame. They consist of the nasion and two pre-auricular points which are digitized during acquisition, and are then also identified by offline visual inspection on the MRI images. Additional digitization points on the head surface can also be used to better adjust the coregistration. In this study, on average, 135 digitization points were available per subject. The transformation, which consists of a rotation matrix and a translation vector, is then typically saved to a small file, also called *trans* file, and later used to compute the forward solution.

For quality insurance, MNE offers a simple function to visualize the result of the coregistration. Figure 9 shows one example obtained with this function with the defaced, low-resolution MRI head surface. As here the MRI were defaced, many important digitization points close to the nose where useless. To reduce the risk of bad coregistration due to defaced MRI images, we used the trans files kindly provided by the original authors.

**Figure 9:**
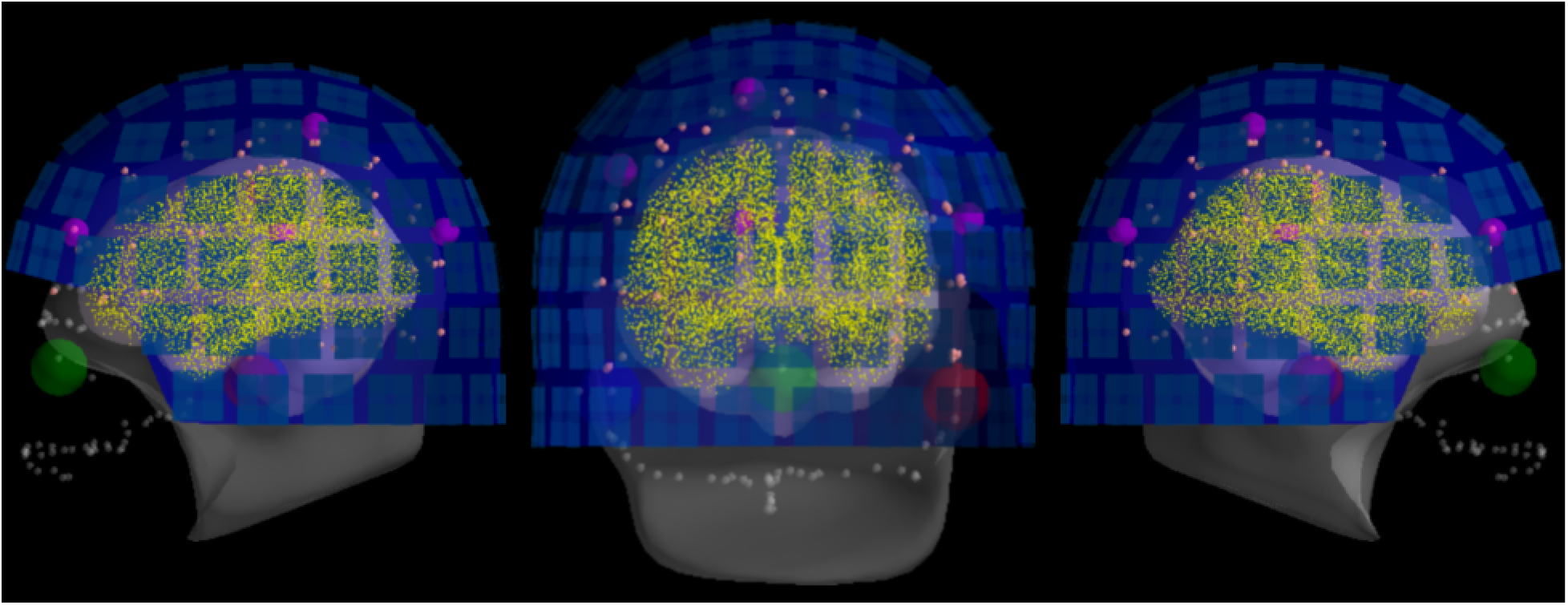
The result of head-to-MRI (and MEG-to-head) transformations with inner skull and outer skin surfaces for one subject. Note that the MEG helmet is well-aligned with the digitization points. The digitized fiducial points are shown with large dots, EEG electrodes with small pink dots, and extra head digitization points with small gray dots. Note that the anonymization of the MRI produces a mismatch between digitized points and outer skin surface at the front of the head.

### 5.4 Covariance estimation and Whitening

As inverse solvers typically assume Gaussian noise distribution on the sensors with an identity covariance matrix, a whitening step is first necessary (Engemann and Gramfort, 2015). M/EEG signals are indeed highly spatially correlated. Whitening also allows integration of data from different channel types that can have different units and signal amplitudes which differ by orders of magnitudes (cf. planar gradiometers, axial magnetometers, and EEG electrodes). To whiten the data, one must provide an estimate of the spatial noise covariance matrix. This can be computed from empty-room recordings for MEG or pre-stimulus periods (Gramfort et al., 2014). Here, we followed the approach proposed by Engemann and Gramfort (2015), which consists in picking the best model and estimating the best regularization parameters by computing the Gaussian log-likelihood of left-out data (i.e., a cross-validation procedure). Such an approach has been shown to be particularly robust for scenarios where a limited number of samples is available for covariance estimation.

In this analysis, the noise covariance is estimated from the 200 ms of data before stimulus presentation. During this period, only a fixation color is visible at the center of the screen. Given this covariance matrix and the gain matrix, one can assemble the inverse operator to compute the MNE or dSPM solutions (Dale et al., 2000a).

The quality of the covariance estimation and whitening can have a significant impact on the source localization results. The rank-adjusted global field power (GFP) has been proposed by Engemann and Gramfort (2015) as a measure that can be used to check the quality of the whitening. It is defined as 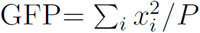 where *P* is the rank of the data and *x_i_* is the signal in the *i*th sensor at a time instant. The GFP being a normalized sum of Gaussian random variables with an identity covariance matrix, it follows a *χ*^2^ distribution with an expected value of 1. What is not captured by our noise model, e.g. actual brain signals, thereof will pop out in the whitened domain. To understand this better, we show some whitened data and the GFP in Figure 10. If the Gaussian assumption has not been violated, we expect the whitened data to contain 95% of the signal within the range of -1.96 and 1.96, which we mark in dotted red lines. The baseline period, where we estimated our noise covariance from, appears to satisfy this assumption. Consequently, the GFP is also 1 during this period. One can observe a strong increase in the GFP just after the stimulus onset, and that it returns slowly to 1 at the end of the time interval. Such a diagnostic plot can in fact be considered essential for quality assurance before computing source estimates. This has as consequence that what appears in the source estimates depends on our noise model. For instance, using a noise covariance obtained from empty room recordings would suggest the presence of “interesting” signals, simply because it contains brain signals that are fundamentally different from the empty room noise.

**Figure 10:**
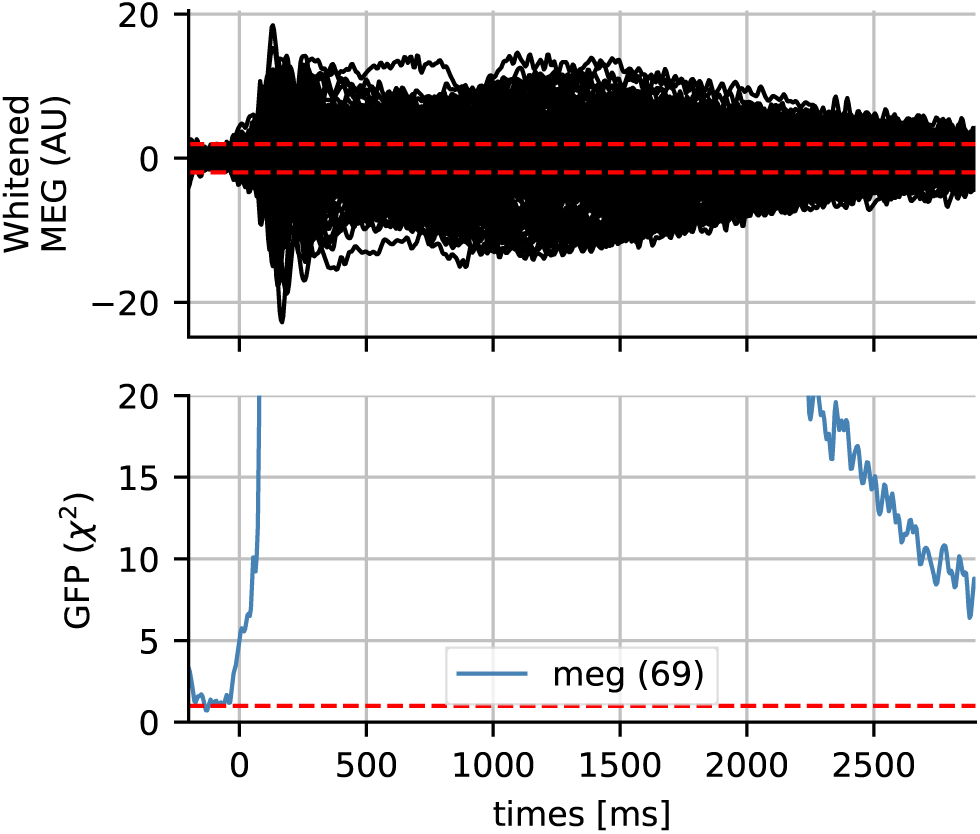
Whitened MEG data for subject 4 and the global field power (GFP) which follows a *χ*^2^ distribution if the data is assumed Gaussian. The dotted horizontal red lines represent the expected GFP during the baseline for Gaussian data. Here the data slowly return to baseline at the end of the epoch.

For the LCMV beamformer, we also need to estimate a signal covariance. For this we use the 30 ms to 300 ms window after the stimulus onset. The data covariance is again regularized automatically following (Engemann and Gramfort, 2015) and is motivated by the results from (Woolrich et al., 2011; Engemann et al., 2015).

#### Caveats

If empty-room data are used to whiten processed signals, one must make sure that the obtained noise covariance matrix corresponds to the processed data rather than to the original empty-room data. This is done by processing the empty-room data with exactly the same algorithm and the same parameters as the actual data to be analyzed. For example if SSS, SSP or ICA are applied on processed data, it should be applied to empty room data before estimating the noise covariance. Concretely, SSP vectors and ICA components projected out from the data of interest should also be projected out from the empty room data. SSS should be performed with identical parameters. Also note that magnetometers and gradiometers are whitened jointly. Moreover, if SSS was applied, the display of whitening treats magnetometers and gradiometers as one channel-type. For proper assessment of whitening, a correct assessment of the spatial degrees of freedom is necessary. The number of SSS dimensions is commonly a good estimate for the degrees of freedom. When movement compensation was applied, the estimated data rank maybe unreliable and suggest too many independent dimensions in the data. Even the actual number of SSS components can be misleading in such circumstances. It is then advisable to inspect the eigen-value spectrum of the covariance matrix manually and specify the degrees of freedom manually using the rank parameter.

### 5.5 Inverse solvers and beamforming

The goal of an inverse solver is to estimate the locations and the time courses of the sources that have produced the data. While the data **M** can be expressed linearly from the sources **X** given the gain matrix **G**, **M ≈ GX**, the problem is ill-posed. Indeed **G** has many more columns than rows. This means that there are more unknown variables (brain sources) than the number of measured values (M/EEG sensors) at each time point. This also implies that the solution of the inverse problem is not unique.

For this reason, many inverse solvers have been proposed in the past ranging from dipole fits (Scherg and Von Cramon, 1985; Mosher et al., 1992), minimum norm estimates (MNE) (Hämäläinen and Ilmoniemi, 1984), and scanning methods such as RAP-MUSIC or beamformers such as LCMV and DICS (Van Veen et al., 1997; Gross et al., 2001; Sekihara et al., 2005). There is therefore no absolute perfect inverse solver, although some are more adapted than others depending on the data. Some are adapted to evoked data for which one can assume a few set of focal sources. Some also give you source amplitudes in a proper unit, which is nAm for electrical current dipoles, such as MNE, MxNE Gramfort et al. (2013b) or dipole fits. Some give you spatially normalized statistical maps such as dSPM (Dale et al., 2000b) or LCMV combined with neural activation index (NAI) filter normalization (Van Veen et al., 1997).

Given the important usage of dSPM and the LCMV beamformer in the cognitive neuroscience literature, we wanted to investigate how much using one of these two most commonly used methods was affecting the source localization results. The dSPM solution was computed with MNE default values: loose orientation of 0.2, depth weighting (Lin et al., 2006) of 0.8, and SNR value of 3. The LCMV used was a vector beamformer with unit-noise-gain normalization (Sekihara et al., 2005) as implemented in MNE 0.15. No specific regularization was used in the beamformer filter estimation.

### 5.6 Group source reconstruction

To analyze data at the group level, some form of data normalization is necessary, whereby data from all subjects is transformed to a common space in a manner that helps compensate for inter-subject differences. This procedure, called *morphing* by the MNE software, exploits the FreeSurfer spherical coordinate system defined for each hemisphere (Dale et al., 1999; Fischl et al., 1999). In our analysis, the data are morphed to the standard FreeSurfer average subject named `fsaverage`. The morphing procedure is performed in three steps. First, the subsampled data defined on the high resolution surface are spread to neighboring vertices using an isotropic diffusion process. Next, registration is used to interpolate the data on the average surface. And finally, the data defined on the average surface is subsampled to yield the same number of source locations in all subjects (here, 10242 locations per hemisphere). Once the morphing is complete, the data is simply averaged.

What is presented in Figure 11 is the group average of the dSPM and LCMV beamformer solutions on contrast between faces and scrambled at 170 ms post-stimulus.

**Figure 11:**
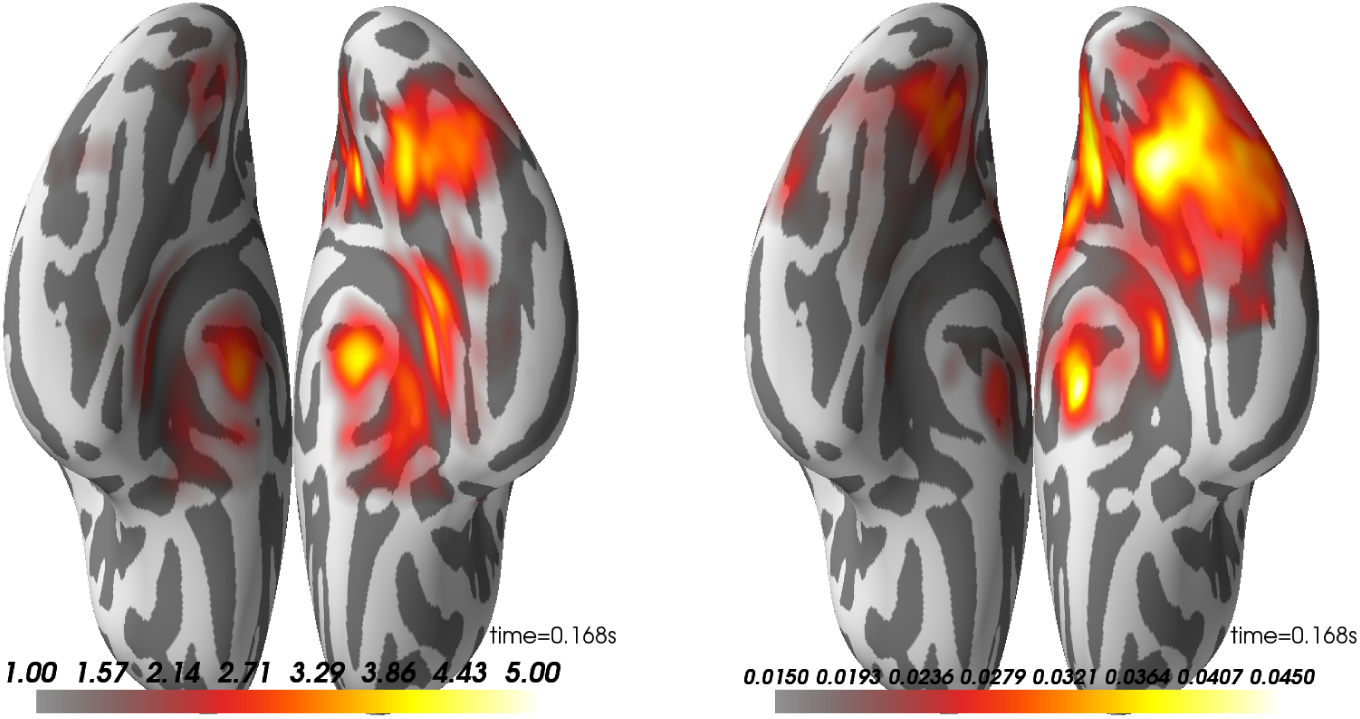
Group average on source reconstruction with dSPM (left) and LCMV (right). Here, we have the ventral view of an inflated surface with the anterior-posterior line going from the bottom to top of the image. Right hemisphere is on the right side.

Looking at these results, one can observe that both methods highlight a peak of activation on the right ventral visual cortex known to be involved in face processing (Grill-Spector et al., 2017, 2004; Wakeman and Henson, 2015). The dSPM peak seems however to be slightly more anterior.

### 5.7 Source-space statistics

Just as we did for the sensor time courses, we can subject the source time courses (here for dSPM only) to a cluster-based permutation test. The null hypothesis is again that there is no significant difference between the data distributions (here measured using cluster size) for faces versus scrambled (paired). Under each permutation, we do a paired t-test across subjects for the difference between the (absolute value of the) faces and scrambled values for each source space vertex and time point. These are clustered, and maximal cluster size for each permutation is selected to form the null distribution. Cluster sizes from the actual data are compared to this null; in this case we find three clusters that lead us to reject the null with *p* < 0.05 (see Figure 12).

**Figure 12:**
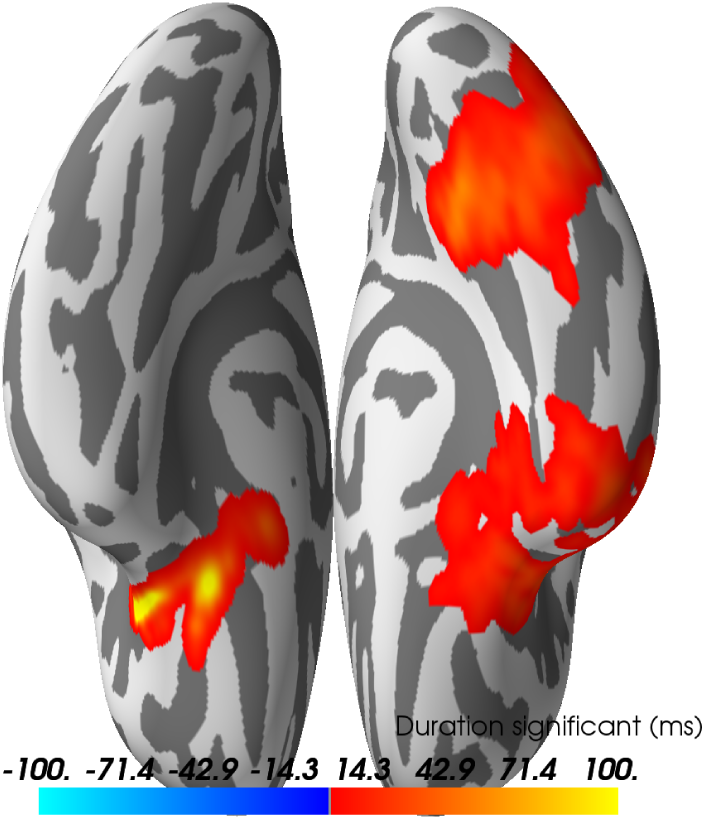
Spatio-temporal source space clusters obtained by nonparametric permutation test that allowed rejection of the null hypothesis that the distribution of data for the “faces” condition was the same as that of “scrambled”. The clusters here are collapsed across time such that vertex colors indicate the duration that each vertex was included in its cluster (each cluster here occurring with FWER corrected *p* < 0.05). Hot colors indicate durations for vertices in clusters where response for faces > scrambled (cool colors would be used for scrambled > faces, but no such clusters were found).

#### Alternatives

When strong hypotheses exist with regard to spatial, temporal and spectral regions of interest, it may be preferable to test the experimental hypotheses on fewer well-chosen signals. In the context of a group analysis, a linear multilevel modeling approach may provide an interesting option for obtaining joint inference at the single subject and group level Gelman (2006); Baayen et al. (2008).

## 6 Discussion and conclusion

Analyzing M/EEG requires successive operations and transformations on the data. At each analysis stage, the different processing choices can affect the final result in different ways. While this situation encourages tailoring data analysis strategies to the specific demands of the scientific problem, this flexibility comes at a cost and can lead to spurious findings when not handled appropriately (Ioannidis, 2005b; Simmons et al., 2011; Carp, 2012a). In the absence of fully automated data analysis pipelines that can optimize the choice of processing steps and parameters, it is crucial to develop principled approaches to planning, conducting and evaluating M/EEG data analysis.

The present study makes the effort to elucidate common elements and pitfalls of M/EEG analysis. It presents a fully reproducible group analysis of the publicly available dataset from Wakeman and Henson (2015). All code and results are publicly accessible http://mne-tools.github.io/mne-biomag-group-demo/. The study provides contextualized in-depth discussion of all major steps of the analysis with regard to alternative options, caveats and quality control measures. As a rare contribution to the M/EEG literature, this study illustrates in comparative figures, the experimental results obtained when changing essential options at different steps in the analysis. In the following, we want to share some insights that we obtained from working together on this study.

### Collaborative data analysis

In our experience, high-level planning and hands-on data analysis are commonly divided between, e.g., masters or doctoral students, post-docs, and senior researchers. As a consequence, the results are typically appreciated from figures produced without connection to the research code that generated them. In this study, several authors contributed repeatedly to the code, analyses were repeated on different computers, and results were inspected in an ongoing fashion by many authors. This experience has had as consequence that incoherences, model violations, and other quality concerns were perhaps detected more often than usual, which has greatly contributed to the overall quality of the data analysis. While it is perhaps too extreme or onerous to recommend adopting social interaction habits from open source software development— such as peer review, pair or extreme programming—in scientific data analysis, we believe that data analysis should not be done in isolation. In order to enable full-blown collaborative data analysis in research, analysis must be repeatable, hence, scripted, and a minimum of code organization and readability must be enforced. On the other hand, the best coding efforts will have limited impact if there are not multiple authors with fresh and active data analysis habits. We hope that the example stated by this paper, together with the open source tools and the community it is built upon, can stimulate more collaborative approaches in M/EEG research.

### The costs of reproducibility

It is a commonly neglected reality that reproducibility comes at a price. Making an analysis strictly reproducible not only requires intensified social interactions, hence more time, but also demands more computational resources. It is a combinatorially hard problem if one were to consider all the potential sources of variability. For example, analyses have to be repeated on different computers with different architectures and performance resources. This sometimes reveals differences in results depending on the hardware, operating system, and software packages used. As observed in the past by Glatard et al. (2015), we noticed that some steps in our pipeline such as ICA are more sensitive to these changes, eventually leading to small differences at the end of the pipeline, which is in our case are cluster-level statistics in the source space. Of course, differences due to these changes are harder to control and enforce in the context of today’s fast technological progress. Indeed, what we manage to achieve is reproducibility, as opposed to the pure replicability which would be the case if the same results could be achieved even when the computer hardware and software packages were changed.

Also, when code is developed on large desktop computers which is common in many laboratory settings, replication efforts with lower-performance workstations may incur high costs in terms of human processing time. The analysis not only runs slower but may crash, for example due to differences in computer memory resources. We therefore emphasize the responsibility of software developers in providing scalable performance control and the responsibility of hands-on data analysts to design the analysis bearing performance and social constraints in mind. In other words, consider that code needs to run on someone else’s computer.

### When to stop?

Obviously, in the light of the current replication crisis, clear rules need to be established on when to stop improving the data analysis (Simmons et al., 2011; Szucs and Ioannidis, 2017). A particular risk is emanating from the possibility of modifying the analysis code to eventually confirm the preferred hypothesis. This would invalidate inference by not acknowledging all the analysis options explored. Apart from commonly recommended preregistration practices and clean hold out data systems, we want to emphasize the importance of quality criteria for developing the analysis. The bulk of M/EEG preprocessing tasks are either implicitly or explicitly model-based, as shown by the rich battery of quality control visualizations presented in this manuscript. Such plots allow to assess if M/EEG analysis outputs can be considered good signals. Consequently, analysis should be stopped when no further improvement on quality control metrics is to be expected, within a reasonable time investment. In other words, not research hypotheses (and statistical significance of results) but rather signal quality metrics are the criterion for constructing M/EEG analyses. Ideally, only when quality control is done, should the contrast(s) of interest be investigated.

With these broader insights in mind, we will make an attempt to extract from our analysis practical recommendations that should facilitate *future* M/EEG analyses. We encourage the reader not to take the analysis presented here as a direct justifications for parameter choices used in their analyses, but instead learn the principles underlying the choices made in our examples. The general rule is: assess your options and chose the optimal measure at each processing step, then visualize and automate as much as you can.

Practical recommendations:

1. **Know your I/O**. Make sure to have a clear idea about the meta-data available in your recordings and that the software package knows about relevant auxiliary channels, e.g, stim, EOG, ECG. Use custom MNE functions and other libraries to add quick reading support if I/O for a file-type is not readily supported.
2. **Think noise**. Inspect your raw data and power spectra to see if and how much denoising is necessary. When using methods such as SSS, SSP, ICA, or reference-channel correction, be aware of their implications for later processing. Remember also to process your empty room data the same way. The interpretation of sensor types may change. Denoising may implicitly act as a high-pass filter (cf. tSSS). High-pass filtering or baselining may not be a good thing, depending on the paradigm. For calibrating your inverse solution, think of what is an appropriate noise model, it may be intrinsically linked to your hypothesis.
3. **Mind signals of non-interest**. Detect and visualize your physiological artifacts, e.g. ECG, EOG, prior to attempting to mitigate them. Choose an option that is precise enough for your data. There is no absolute removal, only changes in signal-to-noise ratio. Not explicitly suppressing any artifacts may also be a viable option in some situations, whereas a downstream method (e.g., temporal decoding) will not benefit from them. When employing an artifact removal technique, visualize how much of your signal of interest is discarded.
4. **Visually inspect at multiple stages**. Use diagnostic visualizations often to get a sense of signal characteristics, from noise sources, to potential signals of interest. Utilize knowledge of paradigms (e.g., existence of an N100 response) to validate steps. Visual inspection of data quality and SNR is recommended even if the processing is automated. When using the an anatomical pipeline, look at your coregistration and head models to make sure they are satisfactory. Small errors can propagate and induce spurious results. Check for model violations when working with inverse solvers and understand them. Inappropriate noise models will distort your estimated sources in simple or complex ways and may give rise to spurious effects.
5. **Apply statistics in a planned way**. Averaging data is a type of statistical transformation. Make sure that what you average is actually comparable. To handle the multiple-comparisons problem, different options exist. Non-parametric hypothesis-tests with clustering and multivariate decoding are two such options, and they are not mutually exclusive. Keep in mind that MEEG is primarily about time, not space. A whole-brain approach may or may not be the best thing to pursue in your situation. Anatomical labels may provide an effective way of reducing the statistical search space.
6. **Be mindful of non-deterministic steps**. To maximize reproducibility, make sure to fix the random initialization of non-deterministic algorithms such as ICA. Not only does it ensure reproducibility, debugging is also easier when the code is deterministic. Prefer automated scripts as opposed to interactive or manual pipelines wherever possible.
7. **Keep software versions fixed**. In an ideal world, software (and hardware) versions would not matter, as each operation necessary for data analysis should be tested against known results to ensure consistency across platforms and versions. However, this ideal cannot always be met in practice. To limit difficulties, do not change software versions, hardware or operating system versions in the middle of your analysis. Keep in mind that MNE is based on several other pieces of software. Updating them can have an impact on the outcome of MNE routines. Once data analysis is complete, cross-checking on different platforms or with different software versions can be useful for community feedback and identifying fragile or problematic steps.

In order to facilitate the reproduction of all the results presented in this manuscript, all the code used to make the figures in this paper, but also much more, is available at http://mne-tools.github.io/mne-biomag-group-demo/.

## Acknowledgement

We would like to thank the many members of the MNE community who have contributed through code, comments, and even complaints, to the improvement and design of this software.

During the preparation of this manuscript, we benefited greatly from discussions with other software package maintainers, such as Brainstorm maintainers who suggested that we visualize PSDs on a channel-by-channel basis to more effectively identify bad sensors. We thank Sami Aboud for discussions on the manuscript and proof-reading.

This work was supported by National Institute of Biomedical Imaging and Bioengineering grants 5R01EB009048, NIH R01 MH106174 and P41RR014075, National Institute on Deafness and Other Communication Disorders fellowship F32DC012456, NSF awards 0958669 and 1042134, the French National Research Agency (ANR-14-NEUC-0002-01) and the European Research Council Starting Grant SLAB ERC-YStG-676943.

## Footnotes

1 http://github.com/mne-tools/mne-biomag-group-demo

2 http://martinos.org/mne/stable/manual/io.html

3 http://martinos.org/mne/stable/auto_tutorials/plot_info.html

4 http://support.hdfgroup.org/HDF5/

5 http://martinos.org/mne/stable/auto_tutorials/plot_artifacts_correction_filtering.html

6 http://martinos.org/mne/stable/auto_tutorials/plot_epoching_and_averaging.html

7 http://martinos.org/mne/stable/auto_tutorials/plot_epoching_and_averaging.html

8 http://martinos.org/mne/stable/auto_tutorials/plot_object_epochs.html

